# Characterization of novel Actinobacteriophage Giantsbane reveals unexpected cluster AU relationships

**DOI:** 10.1101/2020.01.10.891226

**Authors:** Pei Ying Chen, Christopher Liu, Preston Dang, Michael Zhang, Andrew Kapinos, Ryan Ngo, Krisanavane Reddi, Jordan Moberg Parker, Amanda C. Freise

**Author notes:** Current address: Department of Pathology & Laboratory Medicine, Children’s Hospital Los Angeles, Los Angeles, CA, USA. Current address: Department of Pathology, Olive View-UCLA Medical Center, Los Angeles, CA, USA. Current address: Department of Public Health and Community Medicine, Tufts University School of Medicine, Boston, MA, USA. Current address: Thermo Fisher Scientific, Carlsbad, CA, USA. Current address: School of Biological Sciences, Department of Molecular Biology, University of California, San Diego, CA, USA. Current address: School of Medicine, Department of Pediatrics, University of California, San Diego, CA, USA. Current address: College of Dental Medicine, California Northstate University, Elk Grove, CA, USA. Current address: Department of Biomedical Science, Kaiser Permanente Bernard J. Tyson School of Medicine, Pasadena, CA, USA. Equal contributions. Corresponding Author: Amanda C. Freise.

## Abstract

Bacteriophages that infect *Arthrobacter*, a genus of bacteria which play key ecological roles in soil, warrant further study. Giantsbane, a novel Actinobacteriophage, was isolated using *Arthrobacter globiformis* as a host. Transmission electron microscopy and whole-genome sequencing revealed a *Siphoviridae* morphology and a genome length of 56,734 bp. Genome annotation identified 94 putative genes, such as a duplicated major tail protein and a major capsid and protease fusion protein. No genes were associated with lysogeny, indicating a lytic phage. Giantsbane was assigned to the phage cluster AU. Batch average nucleotide identity analysis and phylogenetic networks constructed from shared genes revealed unexpected nucleotide and gene content similarities within cluster AU. These findings have resulted in the creation of two new AU subclusters and the resubclustering of three AU bacteriophages. Analysis using Phamerator and MEME identified repeated motifs and a gene cassette present in all evaluated cluster AU phages which may promote recombination. These findings offer the first intra-cluster analysis of cluster AU phages and further our understanding of the relationships between closely related bacteriophages.

## INTRODUCTION

Bacteriophages (phages), the most abundant biological entities, are still not well-understood or characterized [1]. Soil phages are of particular interest because of their ability to infect bacteria critical to ecological processes such as nutrient cycling [2]. Characterizing these phages’ genomes and the proteins they express increases our understanding of their biological and ecological roles, and contributes to a significant and growing body of knowledge. *Arthrobacter*, a genus of soil bacteria, is a keystone taxon in soil ecosystems due to its ability to recycle nitrogen and carbon [3]. The diverse metabolic capabilities of this genus affect the composition of nutrients in soil, impacting soil biodiversity and health [4]. Understanding the phages that infect this host genus is of particular importance given the key role these bacteria play in the environment.

Phages shape the microbiomes of different environments through phage-host interactions, and frequently exchange genes with each other and their hosts, leading to incredible diversity in the phage population. Organizing, or clustering, these diverse phages allows us to characterize and compare them, giving us more insight into phage evolution and diversity. Clustering methodology is fluid, and changes based on available data. Actinobacteriophages were originally clustered based on genomic nucleotide identity [5]; as more phages were sequenced, these parameters were modified for some hosts. For example, phages from *Gordonia* and *Microbacterium* hosts are clustered based on gene content similarity [6,7]. Nucleotide identity was used as the primary parameter for clustering *Arthrobacter* phages [8]; some new clusters of *Arthrobacter* phages are now created based on gene content similarity [9]. The same generic parameters are used to divide each cluster into subclusters, as phages within the same cluster can be diverse. Inter- and intra-cluster comparative genomic analyses are necessary for more accurate characterization of phages since the present parameters may not fit every cluster. Here we report the isolation, phenotypic characterization, and genomic analysis of a novel cluster AU *Arthrobacter* phage, Giantsbane.

Giantsbane was isolated using the host bacteria *Arthrobacter globiformis* and transmission electron microscopy (TEM) revealed that Giantsbane was a member of the *Siphoviridae* family. Whole-genome sequencing and annotation revealed a genome of 56,734 bp and 94 putative genes. Comparative genomic analyses showed that Giantsbane shares a high nucleotide identity with *Arthrobacter*-infecting cluster AU phages. Further examination revealed unexpected intra-cluster relationships within cluster AU, based on nucleotide and pham similarities, and led to the creation of subclusters AU4 and AU5. A 32 bp repeat sequence found in all cluster AU phages appears to facilitate homologous recombination and may provide a basis for genetic mosaicism, increasing phage diversity. These are the first comparative genomic analyses involving cluster AU phages to date.

## MATERIALS AND METHODS

### Phage isolation

A soil sample collected in Los Angeles, CA (34.065562°N, 118.441146°W) was incubated with enrichment 2X PYCa broth (yeast extract 1 g/L, peptone 15 g/L, 4.5mM CaCl_2_, dextrose 0.1%) at 250 rpm with shaking at 25°C for 1.5 hours and filtered through a 0.22 μm syringe filter. A 2:1 mixture of filtrate and *A. globiformis* strain B-2979 was incubated for 20 minutes before a PYCa double agar overlay and 24-hour incubation.

Spot tests with PYCa and *A. globiformis* B-2979 were performed for select putative plaque clearings [10]. A spot test clearing was picked and diluted in phage buffer (10 mM Tris stock pH 7.5, 10 mM MgSO_4_ stock, 68 mM NaCl, 1 mM CaCl_2_). Two purification plaque assays with *A. globiformis* B-2979 were performed using PYCa double agar overlay, with the modification of a 10-minute absorption before plating [11]. High-titer lysate was collected by flooding webbed-lysis assay plates.

### Transmission electron microscopy

Purified phage lysate was placed on a carbon-coated EM grid for 2 minutes and dried. Lysate was stained with 3 μL of 1% uranyl acetate stain solution for 2 minutes, blotted dry, and washed with ultrapure water. The grid was air-dried for 10 minutes and visualized with a FEI T12 microscope (FEI Company, Hillsboro, U.S.). Measurements of capsids and tails were performed using ImageJ [12].

### DNA extraction, sequencing and assembly

Phage DNA was extracted using the Wizard® DNA Clean-Up System (cat # A7280, Promega, WI, USA). Sequencing libraries were made using a NEBNext® Ultra™ II DNA Library Prep kit (New England Biolabs, MA, USA) and were sequenced using the Illumina MiSeq platform to 2197x coverage. Contigs were assembled using Newbler version 2.9 which was checked for accuracy and genomic termini using Consed version 29 [13].

### Gene location

DNA Master version 5.0.2 (http://cobamide2.bio.pitt.edu/computer.htm) and PECAAN (https://pecaan.kbrinsgd.org/) were used for genome auto-annotation. Glimmer version 3.02 and GeneMark version 2.5 were used to predict open reading frames (ORFs) [14,15]. Gene locations and start sites were corrected using Phamerator and Starterator for manual annotation [16].

### Functional calls

BLASTp, using NCBI and PhagesDB version 2.9.0 databases, was used to assign putative gene function [17–19]. BLAST hits with sequence identities above 35%, query coverage above 75%, and E-values below 1e-7 were prioritized as strong functional evidence. HHpred version 3.2.0 was used to detect structure homology with known proteins in the Protein Data Bank (PDB) [20]. High sequence similarity was preferred, as well as high probability (>80-90%), low E-values (<1e-3), and high coverage (>40-50%). The Conserved Domain Database (CDD) was used to find residue homology with known proteins presented in conserved domain models. TmHmm version 2.0 was used to predict the presence of transmembrane helices from the protein sequence [21]. Only predictions with probabilities over 0.75 were considered.

### Comparative genomic analyses

Sequenced genomes of cluster AU phages used in comparative genomic analyses were obtained from the Actinobacteriophage Database [19]. Batch average nucleotide identity (ANI) analysis was conducted using the command line tool OAU and the USEARCH algorithm version 11 [22]. The resulting OrthoANIu values were visualized by a colored heat map created with Prism version 8 (Graphpad Software, CA, USA).

Shared gene content was evaluated using a network phylogeny. PhamNexus was used to generate a Nexus file containing all phams of each non-draft cluster AU phage. SplitsTree version 4.13.1 was used to produce a phylogenetic network from the Nexus file indicating pham similarity between cluster AU phages.

Repeated sequences were identified in Giantsbane using MEME version 5.1.0 [23]. Phamerator was used to locate the position of the sequences and to identify conserved phams and their order within genomes.

## RESULTS

### Isolation and TEM of *Arthrobacter* phage Giantsbane

Giantsbane was successfully isolated on *A. globiformis* using direct isolation from bulk soil. Spot tests and plaque assays were conducted to obtain a purified phage. Plaque assay plates showed clear plaques averaging 1 mm in diameter (Figure 1a).The majority of *Arthrobacter*-infecting phages are *Siphoviridae* [8]. TEM showed that Giantsbane has a head diameter of 80.7 ± 4.2 nm and a long, flexible tail of length 219 ± 15 nm (Figure 1b). These measurements and morphology are consistent with the characteristics of T5-related *Siphoviridae* [24].

**Figure 1.**
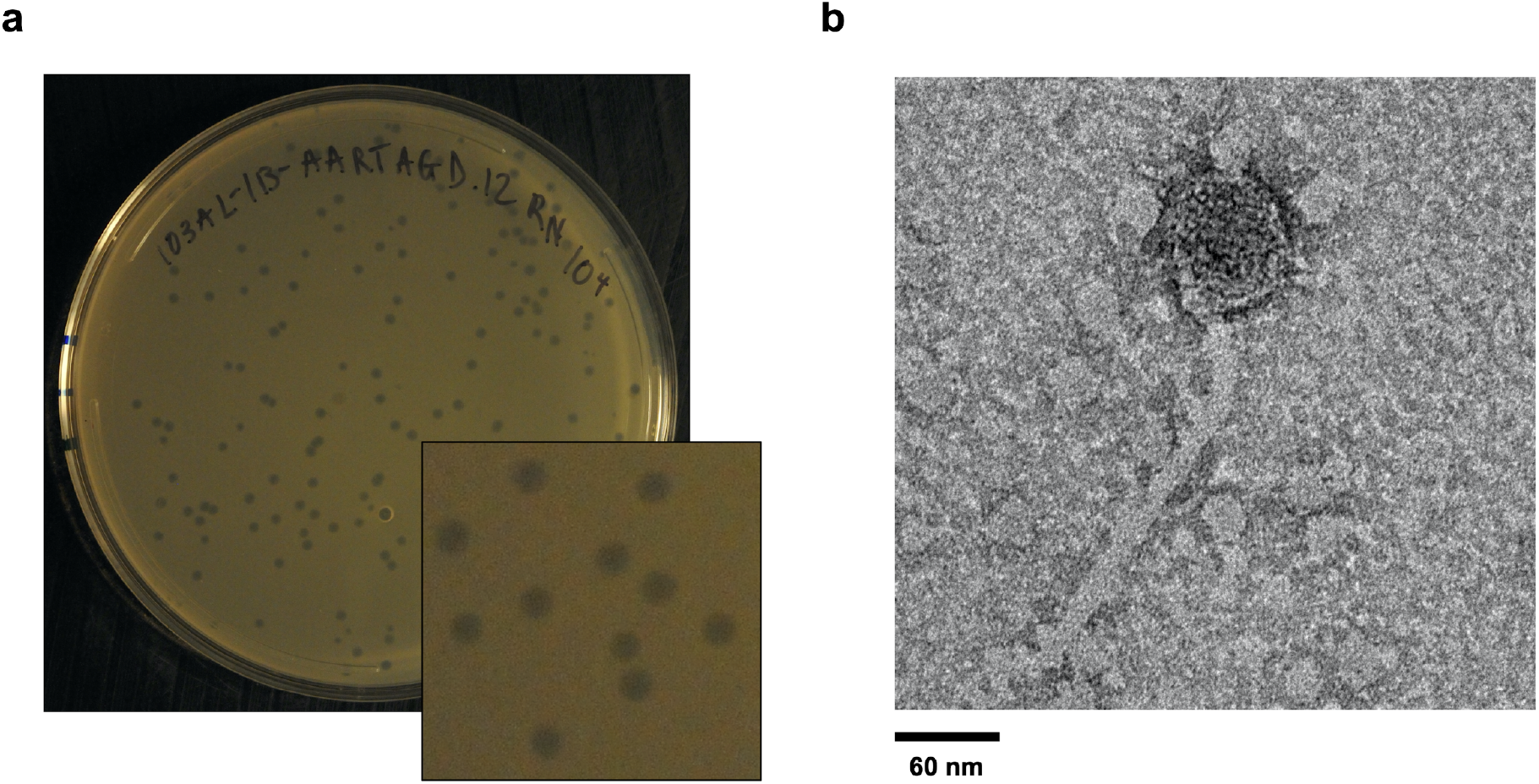
Phenotypic characterization of Giantsbane. **a)** A clearing from the direct isolation plate was purified through two plaque assays to isolate phage Giantsbane. A representative plaque assay plate is shown, with clear plaques averaging 1 mm in diameter. **b)** Transmission electron micrograph of Giantsbane at 52,000x magnification. The diameter of the phage head is 80.7 ± 4.2 nm and the length of the tail is 219 ± 15 nm.

### Giantsbane genome characteristics

The Giantsbane genome was 56,734 bp in length, with a 50.1% GC content. The genome ends had 3’ sticky overhangs, exhibiting a sequence of CGCCGGCCT. Giantsbane was categorized into Actinobacteriophage cluster AU, which contained 17 annotated *Arthrobacter*-infecting phages as of December 2019. Genome annotation identified 94 putative forward genes, 24 of which had putative functions, including core structural, replication, assembly and lysis proteins (Figure 2). No lysogenic proteins were identified, indicating a solely lytic life cycle. Giantsbane’s genome includes two copies of the major tail protein gene, Giantsbane_18 and Giantsbane_23, as well as a major capsid and capsid maturation protease fusion protein encoded in Giantsbane_15 (Figure 2). The fusion protein contains a major capsid protein, capsid maturation protease and scaffolding protein fused into one. It is similar to the capsid and scaffolding fusion protein in *E. coli* HK97 as previously described [25].

**Figure 2.**
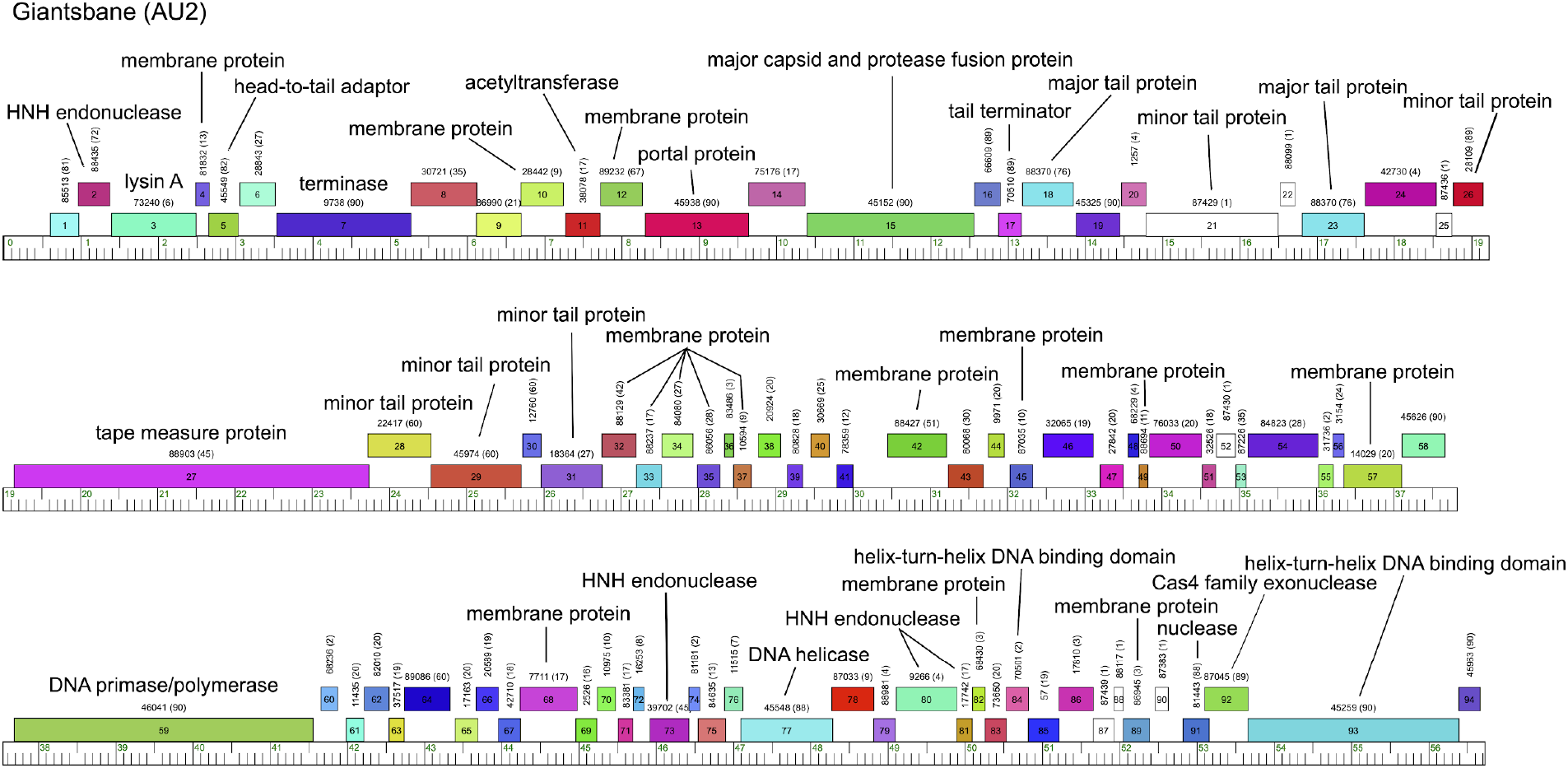
Giantsbane genome map. The phage genome was auto-annotated using DNA Master and then manually corrected with additional genomic analysis programs. The finalized genome contains 94 putative ORFs, all in the forward orientation. 24 ORFs had putative functions assigned to them, including core structural, replication, assembly and lysis proteins.

### Intra-cluster analyses of AU phages reveal unexpected patterns of similarity

Phages are typically sorted into clusters and subclusters to show levels of evolutionary relatedness and facilitate characterization. As of December 2019, cluster AU had 17 annotated members and three subclusters: AU1 (13 phages), AU2 (2 phages), and AU3 (2 phages). Based on nucleotide similarity, Giantsbane was assigned to the *Arthrobacter* phage subcluster AU2. Batch ANI analysis with all non-draft cluster AU phages was used to examine intra-cluster similarities (Figure 3a). All cluster AU pairwise nucleotide identities were above 32.9% of the span-length, with an average of 54.1%. Giantsbane shared the highest percentage nucleotide similarity (86.6%) with AU2 phage Shepard, while AU3 phages Ingrid and Loretta were extremely similar to each other (99.9%). Most of the AU subclusters share 82% or greater gene content within themselves, while different subclusters only share 70-78%. Unexpectedly, phage Makai, which belonged to AU1 at the time of analysis, shared the highest nucleotide similarity with AU2 phages Giantsbane and Shepard (80.5% and 82.2%) out of all cluster AU phages; given its previous assignment to subcluster AU1, Makai was expected to share the highest nucleotide similarity with other AU1 phages. In addition, phages Caterpillar and MediumFry, also previously assigned to AU1, are more similar to one another (94.8%) than to other AU1 phages (77.7%-79.7%). While general guidelines exist for cluster boundaries, subcluster boundaries are less defined. Typically, phages within a subcluster share more genomic similarities with each other than with phages from other subclusters in the same cluster [26].

**Figure 3.**
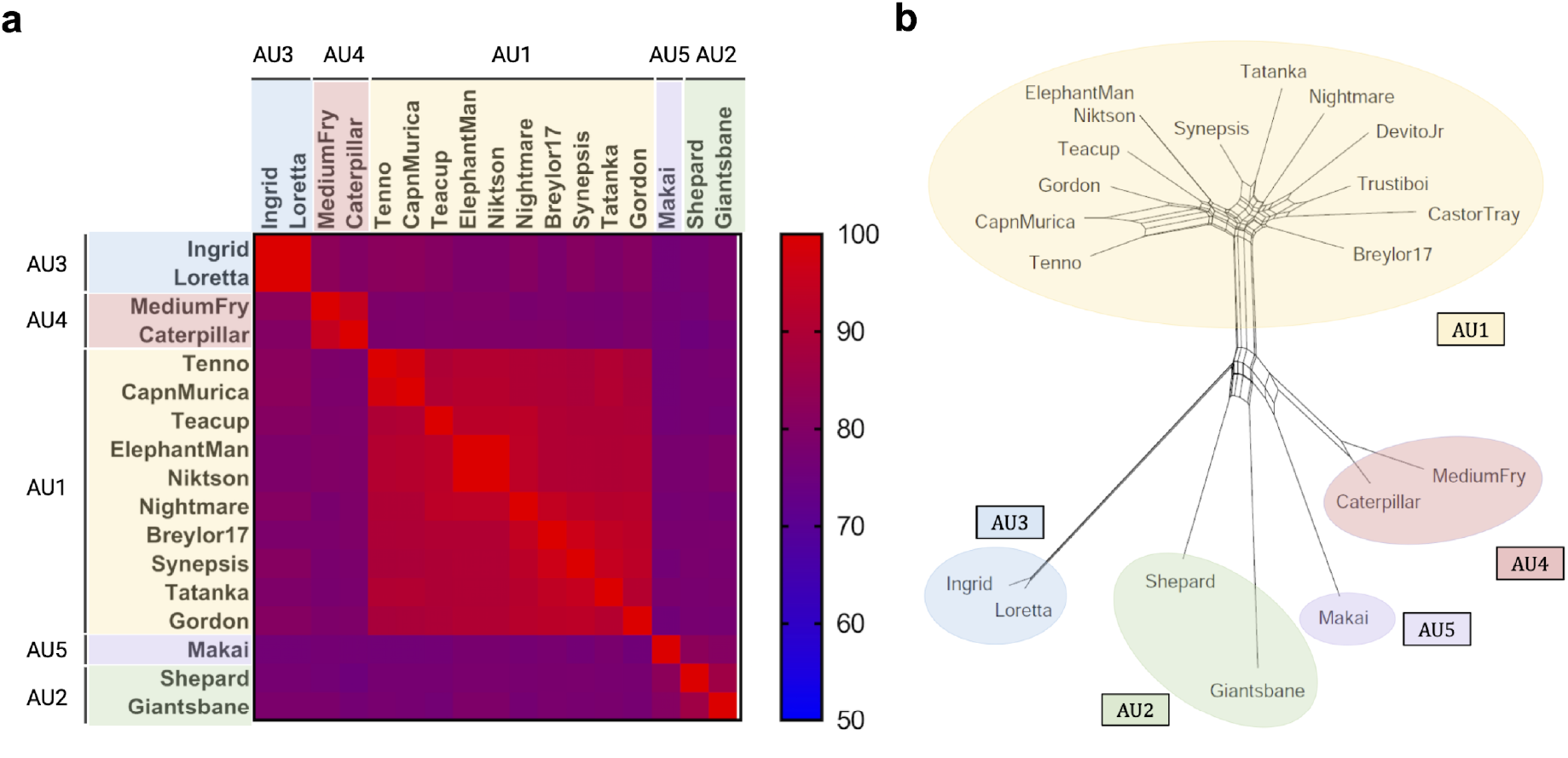
Distinct groupings of AU phages based on nucleotide similarity and shared phams. **a)** A heat map was constructed based on the OrthoANIu values obtained from Batch ANI analysis of 17 annotated cluster AU phages, with a color range of blue to red corresponding to nucleotide percentage similarities of 50% to 100%, respectively. AU5 Caterpillar and MediumFry were shown to be more closely related to each other than to the phages in their previous subcluster, AU1, in terms of nucleotide similarity. **b)** SplitsTree analysis was used to organize 17 annotated and 3 draft cluster AU phages based on pham similarity. Despite their previous AU1 classification, Makai, Caterpillar and MediumFry were shown to be more closely related to the AU2 phages in terms of pham similarity.

At the time of Giantsbane isolation, pham conservation between phages was the secondary parameter for clustering *Arthrobacter* phages. Genes are grouped into phams based on amino acid sequence similarity, and phages sharing at least 35% pham similarity are typically clustered together [26]. Therefore, a phylogenetic network analysis of pham conservation between Giantsbane and other cluster AU phages, visualized through SplitsTree, was used to further examine intra-cluster similarities. Four distinct groups were observed: the first containing Ingrid and Loretta; the second containing Giantsbane and Shepard; the third containing Makai, Caterpillar and MediumFry; and a fourth containing the remaining AU1 phages (Figure 3b). Prior to September 2020, cluster AU consisted of three subclusters, AU1, AU2, and AU3, with phages Caterpillar, MediumFry and Makai belonging to subcluster AU1 (Supplemental Figure 1). Upon examination of these results, together with the ANI analysis, two new subclusters, AU4 and AU5, were created. Based on nucleotide similarity, phages Caterpillar and MediumFry were moved to subcluster AU4 and phage Makai was determined to be the only member of subcluster AU5. These results, together with the ANI analysis, suggest alternative groupings than the established AU subclusters. As of October 2022, three additional AU phages have been isolated and placed in subcluster AU6, as well as an unsubclustered AU phage; these phages were outside the scope of our analysis.

### Cluster AU phages contain repeated sequences flanking conserved phams

Nucleotide motifs, or repeat sequences, may indicate recombination events and drive mosaicism in phages [27,28]. We noticed a cassette of genes in multiple AU phages that appeared to commonly have rearrangements, and decided to examine the genome for potential motifs involved in recombination. Using MEME to search for motifs within a single genome and Phamerator to look at conservation of motifs across genomes, a 32 bp repeat sequence was identified six times in the intergenic regions surrounding genes 37-41 of Giantsbane (Figure 4a). The sequence is indicated by red lines within Phamerator, which correspond to BLASTn alignments with an E-value between 1e-4 and 1e-10 [16]. It occurs immediately upstream and downstream of each gene in this region, making it a boundary sequence as described previously [29]; this pattern is found in all cluster AU phages (Figure 4b, 4d). Figures 4b and 4d show percent nucleotide identity between Giantsbane and representative cluster AU phages in the four cases described below. Genes of this five-gene cassette were subject to deletions and translocations to different regions within the cassette. Case I, shown by Giantsbane and Caterpillar, shows one translocation and one deletion. Case II describes complete pham conservation with one translocation, shown between Giantsbane and Makai (Figure 4c). Case III describes two deletions, shown between Giantsbane and CapnMurica, and Case IV describes three deletions and two translocations, shown between Giantsbane and Ingrid (Figure 4e). Figures 4c and 4e show pham conservation or deletion between Giantsbane and representative cluster AU phages. This cassette contained the only instances of gene translocation observed among cluster AU genomes. Cluster AU6 was not evaluated.

**Figure 4.**
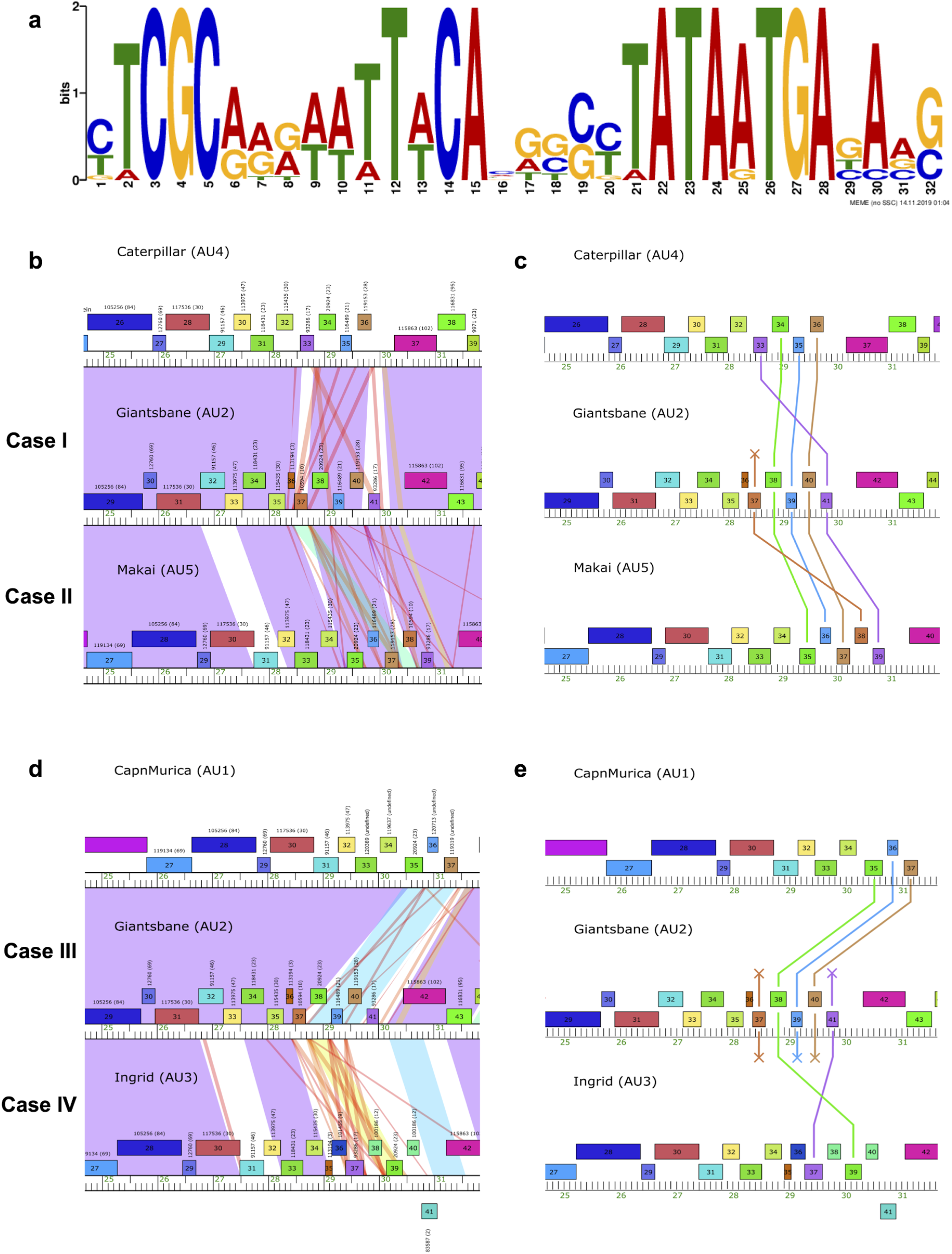
Repeated sequence conserved among AU phages. AU phage genomes were analyzed using MEME and Phamerator to identify repeat sequences, their conservation across different phage genomes, and location relative to genes. **a)** A 32 bp repeat was identified with MEME and displayed as a logogram that shows the likelihood of finding a nucleotide at that position in the sequence. **b)** Phamerator map of Case I and II rearrangements showing nucleotide sequence similarity as shaded areas between representative phage genomes. Purple and red areas indicate high and low E-value BLASTn alignments, respectively, while white areas indicate no similarity. Giantsbane shows high sequence similarity with Caterpillar and Makai in the 28.5–30 kbp region and the red lines correspond to instances of the 32 bp repeat sequence. **c)** PECAAN output showing pham conservation in Case I and II rearrangements. Lines connecting phams between genomes represent pham conservation and lines ending in crosses represent deletions. **d)** Phamerator map of Case III and IV rearrangements with representative phage genomes. Giantsbane shows less sequence similarity with CapnMurica and Ingrid in the 28.5–30 kbp region but the 32 bp repeat sequences, shown by the red lines, are still present. **e)** PECAAN output of Case III and IV rearrangements.

## DISCUSSION

The aim of this study was to investigate the genomic characteristics of novel Actinobacteriophage Giantsbane. Phage biology is a relatively new field; better understanding of phage diversity requires further characterization of novel bacteriophages [1]. Numerous characterizations of *Arthrobacter*-infecting phages exist in the literature, but few include comparative genomics. *Arthrobacter* phages span 30 clusters and multiple singletons as of October 2022, and their genomes currently range from 14,830 bp to 176,888 bp [30]. The majority of previously characterized *Arthrobacter* phages have *Siphoviridae* or *Myoviridae* morphology, but several *Podoviridae* phages have also been documented [31]. The genomes of several phages in cluster AU have been published [32] and cluster AU has been compared to other *Arthrobacter*-infecting clusters [8], but comparative genomic analysis within cluster AU has not yet been reported. Our results add to pre-existing knowledge of *Siphoviridae* and *Arthrobacter*-infecting phages and introduce the first intra-cluster analysis of cluster AU phages to date.

Phages are clustered based on nucleotide and pham similarity to establish evolutionary relationships and facilitate characterization. Batch ANI and pham similarity phylogenetic network analyses revealed that Giantsbane shares over 75% nucleotide similarity over an average span-length of 48% and over 65% pham similarity with other cluster AU phages, confirming Giantsbane’s cluster AU membership. Giantsbane’s lack of lysogenic and tail sheath proteins, as well as its *Siphoviridae* classification based on TEM, are consistent with all Cluster AU members. Giantsbane and other cluster AU phages share many genomic characteristics with other *Arthrobacter*-infecting phages, such as the order of the structural genes — terminase, portal, capsid maturation protease, scaffolding protein, major capsid protein, major tail subunit, tape measure protein and minor tail proteins [8]. Unlike most other *Arthrobacter*-infecting phages, Cluster AU genomes have the lysis cassette upstream of the terminase protein at the left end of the genome and also possess a series of small genes, most of which are putative membrane proteins, downstream of the minor tail proteins [8]. Interestingly, the duplicated major tail protein and the major capsid and protease fusion protein found in Giantsbane are present in all other cluster AU phages, as well as cluster AM, BI, DJ and CC phages, which infect *Arthrobacter*, *Streptomyces*, *Gordonia* and *Rhodococcus*, respectively [32]. The conservation of these proteins across phages infecting different bacterial hosts demonstrates phage genomic mosaicism and warrants further study.

Nucleotide and pham similarity analyses revealed intra-cluster patterns. Cluster AU phages that shared the most similarities were not always subclustered together. Furthermore, subcluster AU1 split into two groups with distinct nucleotide and pham similarities, one consisting of phages Makai, Caterpillar and MediumFry and the other of the remaining AU1 phages. This division within a subcluster has also been observed in *Mycobacterium*-infecting subcluster A3 phages [33]. These inter-subcluster similarities, and the re-categorization of three previous AU1 bacteriophages into newly-created subclusters AU4 and AU5, suggest that subclustering parameters need to be re-evaluated as more phages are discovered. Cases such as singleton Actinobacteriophage BlueFeather’s reclustering into cluster FE, based on shared gene content analysis, further confirm the need for inter- and intra-cluster comparative analyses [9]. These results showcase the diversity and relatedness of *Arthrobacter*-infecting *Siphoviridae* phages, which can help inform and clarify classification of similar phages in the future.

MEME and Phamerator revealed a 32 bp repeat sequence in the intergenic regions of a five-gene cluster in cluster AU phages within the 28.5–30 kbp region. Comparative genomic analyses demonstrated that genes within this region were prone to insertions, deletions or translocations between phages. Several lambdoid phages have shown similar boundary sequences, proposed to create highly mosaic genomic regions [29]. Gene translocations and deletions were only seen in this region in all of the cluster AU genomes, suggesting the potential function of these intergenic repeat sequences. This provides evidence that boundary sequences may provide an additional means of recombination, in addition to illegitimate recombination [34]. Multiple cycles of Giantsbane propagation followed by sequencing of the final generation and comparison with the original genome may help elucidate the importance of intergenic repeat sequences in mechanisms of phage genomic rearrangement and evolution. These findings would further our understanding of the roles of repeat sequences in phages and novel methods of propagating phage diversity.

Nucleotide similarity, pham conservation and repeat sequence analyses revealed complex relationships in the first intra-cluster analysis of cluster AU phages to date. Two new subclusters, AU4 and AU5, were created for three Cluster AU phages and a repeat sequence that may promote recombination was discovered. Further analyses of *Arthrobacter*-infecting phages can lead to better physiological and genomic characterization, as well as a better understanding of phage diversity.

## SUPPLEMENTAL FIGURES

**Figure S1.**
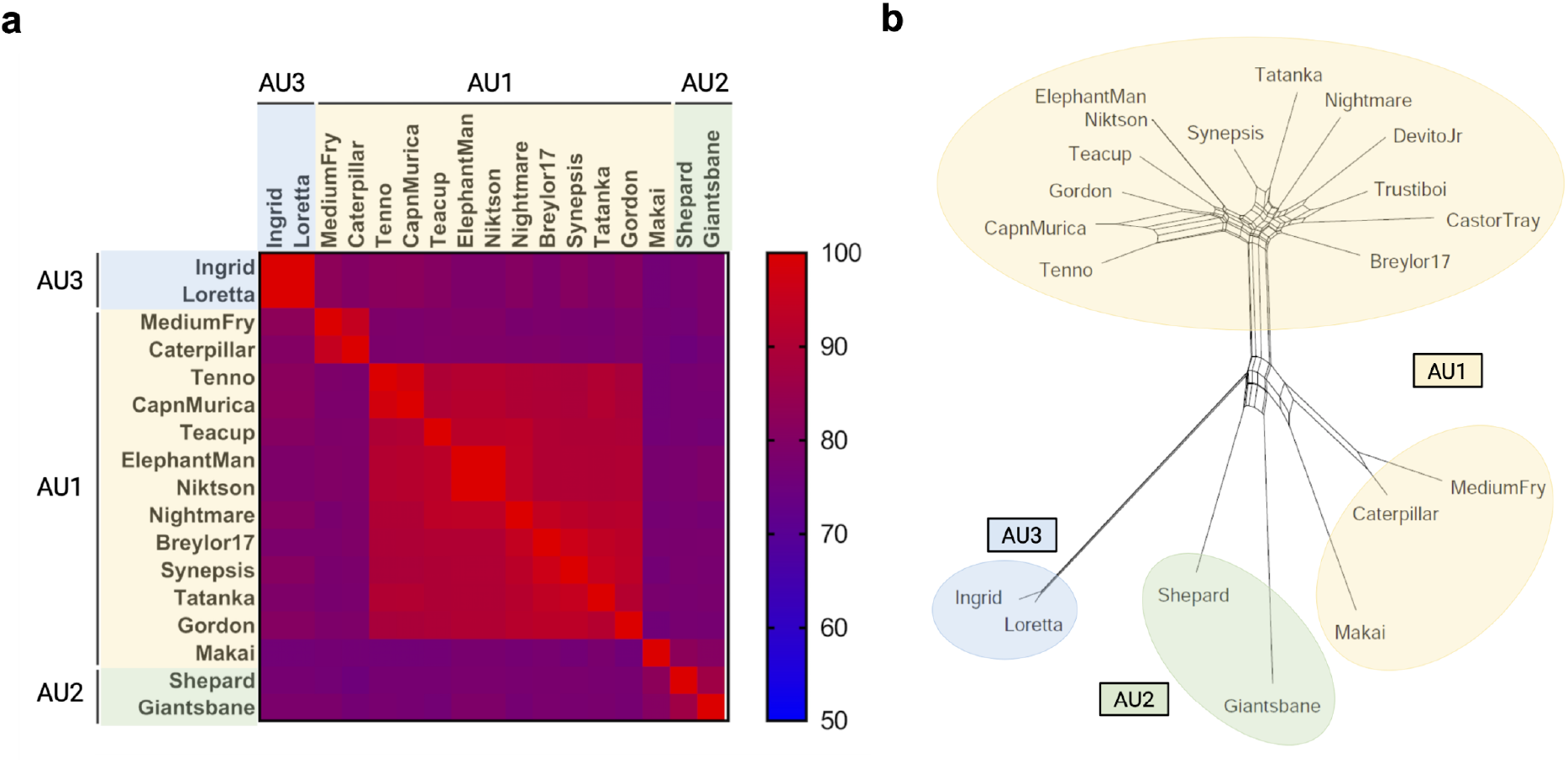
Nucleotide similarity and shared phams analyses of AU phages with original subcluster assignments. **a)** A heat map was constructed based on the OrthoANIu values obtained from Batch ANI analysis of 17 annotated cluster AU phages, with a color range of blue to red corresponding to nucleotide percentage similarities of 50% to 100%, respectively. Caterpillar and MediumFry were shown to be more closely related to each other than to the phages in their previous subcluster, AU1, in terms of nucleotide similarity. **b)** SplitsTree analysis was used to organize 17 annotated and 3 draft cluster AU phages based on pham similarity. Despite their previous AU1 classification, Makai, Caterpillar and MediumFry were shown to be more closely related to the AU2 phages in terms of pham similarity.

## ACKNOWLEDGMENTS

We thank Andy Alag for isolating Giantsbane; Yvette Lakkis and Andrew Lee for laboratory assistance; Rebecca A. Garlena and Daniel A. Russell at the Pittsburgh Bacteriophage Institute for genome sequencing and assembly; and Debbie Jacobs-Sera, Welkin Pope, Graham Hatfull, with the HHMI Science Education Alliance-Phage Hunters Advancing Genomics and Evolutionary Science (SEA-PHAGES) program for programmatic support. We acknowledge the use of instruments at the Electron Imaging Center for NanoMachines supported by NIH (1S10RR23057 to ZHZ) and CNSI at UCLA.

## AUTHORSHIP CONTRIBUTION STATEMENT

P.Y.C., C.L., P.D., and M.Z. drafted the paper and performed experiments; P.Y.C., A.F., and J.M.P. revised the paper; J.M.P. and A.F. supervised the research and performed quality control on the annotations; and all authors contributed to the isolation, annotation, and genome analysis of the novel phages. All authors have reviewed and approved of this manuscript prior to submission.

This project was supported by the Microbiology, Immunology & Molecular Genetics Department and the Dean of Life Sciences Division at UCLA, with additional support for sequencing from the HHMI Science Education Alliance-Phage Hunters Advancing Genomics and Evolutionary Science (SEA-PHAGES) program.

## AUTHORSHIP DISCLOSURE STATEMENT

The authors declare that there is no conflict of interest regarding the publication of this article.

## REFERENCES

1. Hatfull GF. Dark Matter of the Biosphere: the Amazing World of Bacteriophage Diversity. Goodrum F, editor. J Virol. 2015;89: 8107–8110. doi:10.1128/JVI.01340-15

2. Uroz S, Oger P, Lepleux C, Collignon C, Frey-Klett P, Turpault M-P. Bacterial weathering and its contribution to nutrient cycling in temperate forest ecosystems. Res Microbiol. 2011;162: 820–831. doi:10.1016/j.resmic.2011.01.013

3. Romaniuk K, Golec P, Dziewit L. Insight Into the Diversity and Possible Role of Plasmids in the Adaptation of Psychrotolerant and Metalotolerant Arthrobacter spp. to Extreme Antarctic Environments. Front Microbiol. 2018;9: 3144. doi:10.3389/fmicb.2018.03144

4. Eschbach M, Möbitz H, Rompf A, Jahn D. Members of the genus Arthrobacter grow anaerobically using nitrate ammonification and fermentative processes: anaerobic adaptation of aerobic bacteria abundant in soil. FEMS Microbiol Lett. 2003;223: 227–230. doi:10.1016/S0378-1097(03)00383-5

5. Hatfull GF, Jacobs-Sera D, Lawrence JG, Pope WH, Russell DA, Ko C-C, et al. Comparative Genomic Analysis of 60 Mycobacteriophage Genomes: Genome Clustering, Gene Acquisition, and Gene Size. Journal of Molecular Biology. 2010;397: 119–143. doi:10.1016/j.jmb.2010.01.011

6. Jacobs-Sera D, Abad LA, Alvey RM, Anders KR, Aull HG, Bhalla SS, et al. Genomic diversity of bacteriophages infecting Microbacterium spp. Cloeckaert A, editor. PLoS ONE. 2020;15: e0234636. doi:10.1371/journal.pone.0234636

7. Pope WH, Mavrich TN, Garlena RA, Guerrero-Bustamante CA, Jacobs-Sera D, Montgomery MT, et al. Bacteriophages of *Gordonia* spp. Display a Spectrum of Diversity and Genetic Relationships. Losick R, editor. mBio. 2017;8: e01069–17, /mbio/8/4/e01069-17.atom. doi:10.1128/mBio.01069-17

8. Klyczek KK, Bonilla JA, Jacobs-Sera D, Adair TL, Afram P, Allen KG, et al. Tales of diversity: Genomic and morphological characteristics of forty-six Arthrobacter phages. Schuch R, editor. PLoS ONE. 2017;12: e0180517. doi:10.1371/journal.pone.0180517

9. Demo S, Kapinos A, Bernardino A, Guardino K, Hobbs B, Hoh K, et al. BlueFeather, the singleton that wasn’t: Shared gene content analysis supports expansion of Arthrobacter phage cluster FE. PLOS ONE. 2021;16. doi:https://doi.org/10.1371/journal.pone.0248418

10. Kutter E. Phage Host Range and Efficiency of Plating. In: Clokie MRJ, Kropinski AM, editors. Bacteriophages. Totowa, NJ: Humana Press; 2009. pp. 141–149. doi:10.1007/978-1-60327-164-6_14

11. Kropinski AM, Mazzocco A, Waddell TE, Lingohr E, Johnson RP. Enumeration of Bacteriophages by Double Agar Overlay Plaque Assay. In: Clokie MRJ, Kropinski AM, editors. Bacteriophages. Totowa, NJ: Humana Press; 2009. pp. 69–76. doi:10.1007/978-1-60327-164-6_7

12. Abràmoff MD, Magalhães PJ, Ram SJ. Image processing with ImageJ. Biophotonics international. 2004;11: 36–43.

13. Russell DA. Sequencing, Assembling, and Finishing Complete Bacteriophage Genomics. In: Clokie MRJ, Kropinski AM, Lavigne R, editors. Bacteriophages. New York, NY: Humana Press; 2018. pp. 109–125. Available: https://doi.org/10.1007/978-1-4939-7343-9_9

14. Delcher AL, Bratke KA, Powers EC, Salzberg SL. Identifying bacterial genes and endosymbiont DNA with Glimmer. Bioinformatics. 2007;23: 673–679. doi:10.1093/bioinformatics/btm009

15. Besemer J, Borodovsky M. GeneMark: web software for gene finding in prokaryotes, eukaryotes and viruses. Nucleic Acids Research. 2005;33: W451–W454. doi:10.1093/nar/gki487

16. Cresawn SG, Bogel M, Day N, Jacobs-Sera D, Hendrix RW, Hatfull GF. Phamerator: a bioinformatic tool for comparative bacteriophage genomics. BMC Bioinformatics. 2011;12: 395. doi:10.1186/1471-2105-12-395

17. Altschul SF, Gish W, Miller W, Myers EW, Lipman DJ. Basic local alignment search tool. J Mol Biol. 1990;215: 403–410. doi:10.1016/S0022-2836(05)80360-2

18. Wheeler D, Bhagwat M. BLAST QuickStart: example-driven web-based BLAST tutorial. In: Bergman NH, editor. Comparative Genomics. Humana Press; 2007. pp. 149–175. Available: https://doi.org/10.1007/978-1-59745-514-5_9

19. Russell DA, Hatfull GF. PhagesDB: the actinobacteriophage database. Wren J, editor. Bioinformatics. 2017;33: 784–786. doi:10.1093/bioinformatics/btw711

20. Soding J, Biegert A, Lupas AN. The HHpred interactive server for protein homology detection and structure prediction. Nucleic Acids Research. 2005;33: W244–W248. doi:10.1093/nar/gki408

21. Krogh A, Larsson B, von Heijne G, Sonnhammer EL. Predicting transmembrane protein topology with a hidden Markov model: application to complete genomes. J Mol Biol. 2001;305: 567–580. doi:10.1006/jmbi.2000.4315

22. Lee I, Ouk Kim Y, Park S-C, Chun J. OrthoANI: An improved algorithm and software for calculating average nucleotide identity. International Journal of Systematic and Evolutionary Microbiology. 2016;66: 1100–1103. doi:10.1099/ijsem.0.000760

23. Bailey TL, Boden M, Buske FA, Frith M, Grant CE, Clementi L, et al. MEME SUITE: tools for motif discovery and searching. Nucleic Acids Research. 2009;37: W202–W208. doi:10.1093/nar/gkp335

24. Carstens EB. Introduction to Virus Taxonomy. In: King AMQ, Adams MJ, Carstens EB, Lefkowitz EJ, editors. Virus Taxonomy. San Diego, CA: Elsevier; 2012. pp. 1, 3–20. Available: https://doi.org/10.1016/B978-0-12-384684-6.00114-2

25. Veesler D, Khayat R, Krishnamurthy S, Snijder J, Huang RK, Heck AJR, et al. Architecture of a dsDNA Viral Capsid in Complex with Its Maturation Protease. Structure. 2014;22: 230–237. doi:10.1016/j.str.2013.11.007

26. Hatfull GF. Mycobacteriophages. Microbiology Spectrum. 2018;6. doi:10.1128/microbiolspec.GPP3-0026-2018

27. Steiner WW, Steiner EM, Girvin AR, Plewik LE. Novel Nucleotide Sequence Motifs That Produce Hotspots of Meiotic Recombination in *Schizosaccharomyces pombe*. Genetics. 2009;182: 459–469. doi:10.1534/genetics.109.101253

28. De Paepe M, Hutinet G, Son O, Amarir-Bouhram J, Schbath S, Petit M-A. Temperate Phages Acquire DNA from Defective Prophages by Relaxed Homologous Recombination: The Role of Rad52-Like Recombinases. Casadesús J, editor. PLoS Genet. 2014;10: e1004181. doi:10.1371/journal.pgen.1004181

29. Clark AJ, Inwood W, Cloutier T, Dhillon TS. Nucleotide sequence of coliphage HK620 and the evolution of lambdoid phages. J Mol Biol. 2001;311: 657–679. doi:10.1006/jmbi.2001.4868

30. Russell DA, Hatfull GF. PhagesDB: the actinobacteriophage database. Wren J, editor. Bioinformatics. 2017;33: 784–786. doi:10.1093/bioinformatics/btw711

31. Klyczek KK, Jacobs-Sera D, Adair TL, Adams SD, Ball SL, Benjamin RC, et al. Complete Genome Sequences of 44 *Arthrobacter* Phages. Genome Announc. 2018;6: e01474–17, /ga/6/5/e01474-17.atom. doi:10.1128/genomeA.01474-17

32. Adair TL, Stowe E, Pizzorno MC, Krukonis G, Harrison M, Cresawn SG, et al. Genome Sequences of Three Cluster AU Arthrobacter Phages, Caterpillar, Nightmare, and Teacup. Genome Announc. 2017;5: e01121–17, e01121-17. doi:10.1128/genomeA.01121-17

33. Pope WH, Bowman CA, Russell DA, Jacobs-Sera D, Asai DJ, Cresawn SG, et al. Whole genome comparison of a large collection of mycobacteriophages reveals a continuum of phage genetic diversity. eLife. 2015;4: e06416. doi:10.7554/eLife.06416

34. Morris P, Marinelli LJ, Jacobs-Sera D, Hendrix RW, Hatfull GF. Genomic Characterization of Mycobacteriophage Giles: Evidence for Phage Acquisition of Host DNA by Illegitimate Recombination. JB. 2008;190: 2172–2182. doi:10.1128/JB.01657-07

